# Dynamin-2 regulates synaptic podosome maturation to facilitate neuromuscular junction development

**DOI:** 10.1101/2020.05.30.125062

**Authors:** Shan-Shan Lin, Tsung-Lin Hsieh, Gunn-Guang Liou, Tsai-Ning Li, Hsin-Chieh Lin, Chiung-Wen Chang, Hsiang-Yi Wu, Chi-Kuang Yao, Ya-Wen Liu

## Abstract

Neuromuscular junctions (NMJs) govern rapid and efficient neuronal communication with muscle cells, which relies on the proper architecture of specialized postsynaptic compartments. However, the intrinsic mechanism in muscle cells contributing to elaborate NMJ development has been unclear. In this study, we reveal that the GTPase dynamin-2 (Dyn2), best-known for catalyzing synaptic vesicle endocytosis at the presynaptic membrane, is also involved in postsynaptic morphogenesis. We demonstrate that Dyn2 is enriched in the postsynaptic membrane of muscle cells and is involved in the maturation of neurotransmitter receptor clusters via its actin bundling ability. Dyn2 functions as a molecular girdle to regulate synaptic podosome turnover and promote morphogenesis of the postsynaptic apparatus. In *Drosophila* NMJs, Dyn2 is required to organize the postsynaptic actin cytoskeleton and to mediate its electrophysiological activities. Mechanistically, the actin binding, self-assembly, GTP hydrolysis ability, and Y597 phosphorylation of Dyn2 all regulate its actin bundling activity. Together, our study uncovers a role for Dyn2 in cytoskeleton remodeling and organization at the postsynaptic membrane of NMJs.

## Introduction

Neuromuscular junctions (NMJs) control the final output of the nervous system to direct voluntary movements. NMJs are equipped with elaborate membranous and cytoskeletal structures developed from a stepwise process of morphogenesis, i.e., prepatterning of neurotransmitter receptors during embryogenesis and maturation of the pre- and post-synaptic apparatus postnatally (Sanes & Lichtman, 2001). Muscle prepatterning occurs from embryonic day (E) 12.5 to E14.5 of early mouse embryogenesis, which features acetylcholine receptor (AChR) clustering and subsequent motor neuron innervation (Lin, Burgess et al., 2001, Shi, Fu et al., 2012, Yang, Arber et al., 2001). After birth, these AChR clusters undergo a plague-to-pretzel morphological transition that requires extracellular matrix (ECM) signaling, motor neuron activity, and muscle-intrinsic machineries (Bernadzki, Rojek et al., 2014, Bezakova & Ruegg, 2003, Marques, Conchello et al., 2000, Shi et al., 2012). After maturation, NMJs are maintained to enable life-long motor performance with limited plasticity (Burden, Huijbers et al., 2018, Kummer, Misgeld et al., 2006, Shi et al., 2012). Among these processes, the intrinsic mechanism in muscle cells that contributes to NMJ maturation remains poorly understood.

Several studies have identified the presence of a unique actin-based structure, the podosome, at the postsynaptic membrane of NMJs, which dictates the plague-to-pretzel morphological transition of AChR clusters in aneurally-cultured myotubes (Chan, Kwan et al., 2020, Proszynski, Gingras et al., 2009). Podosomes are protrusive actin structures equipped with ECM degradative abilities and they are also involved in proper tissue development, immune cell patrolling, and cancer cell metastasis (Linder, 2007, Linder & Aepfelbacher, 2003, Linder & Wiesner, 2016, Luxenburg, Geblinger et al., 2007). Unlike the podosomes in motile cells that facilitate cell motility, synaptic podosomes promote NMJ maturation by redistributing AChR and ECM components in cultured myotubes (Bernadzki et al., 2014, Kishi, Kummer et al., 2005, Proszynski & Sanes, 2013). Recently, activity of the podosome component MT1-MMP (a metalloproteinase) has been reported to be crucial for NMJ development in *Xenopus* and mouse (Chan et al., 2020). Therefore, although it has been well documented that synaptic podosomes play critical roles in NMJ development, their dynamics and architecture remain ill-defined.

Previously, we discovered that a GTPase, dynamin-2 (Dyn2), is enriched at podosomes in differentiated myoblasts where it stiffens the podosomes and promotes myoblast fusion. Dynamin is a mechanochemical enzyme, most studied for its roles in catalyzing membrane fission during endocytosis and synaptic vesicle recycling (Antonny, Burd et al., 2016, Ferguson & De Camilli, 2012, Hinshaw, 2000). Although Dyn2 is a ubiquitously expressed dynamin isoform, mutations of Dyn2 have been linked to two tissue-specific autosomal-dominant congenital diseases, i.e., Charcot-Marie-Tooth (CMT) neuropathy and centronuclear myopathy (CNM) (Bitoun, Maugenre et al., 2005, Romero, 2010, Zhao, Maani et al., 2018, Zuchner, Noureddine et al., 2005). The CMT-associated Dyn2 mutant proteins are hypoactive and cause defects in endocytosis and myelination of peripheral neurons, whereas CNM-associated Dyn2 mutant proteins are hyperactive and elicit fragmentation of muscular plasma membrane invagination in *Drosophila* and defective triad structures of the skeletal muscle in mouse (Chin, Lee et al., 2015, Cowling, Toussaint et al., 2011, Gibbs, Davidson et al., 2014, Sidiropoulos, Miehe et al., 2012).

Apart from its functions in catalyzing membrane fission, Dyn2 has also been found to localize and function as an actin remodeling protein at many actin-rich structures, including lamellipodia, dorsal membrane ruffles and podosomes (Ferguson & De Camilli, 2012, Menon & Schafer, 2013). Although it has been reported that Dyn2 can regulate actin rearrangement by directly binding actin filaments and interacting with actin polymerization regulators (Gu, Yaddanapudi et al., 2010, Mooren, Kotova et al., 2009), it is not clear what the molecular function of dynamin is at the synaptic podosomes of myotubes.

In this study, we reveal that Dyn2 forms a belt-like structure around the podosome core of myotubes through its actin bundling activity to promote the maturation and turnover of synaptic podosomes, thereby regulating the development and function of NMJs. The actin bundling activity of Dyn2 is derived from its actin-binding and self-assembly abilities, and is regulated by phosphorylation of residue Y597 and GTP hydrolysis. Together, our results reveal a pivotal role for Dyn2 in synaptic podosome maturation and turnover, providing important insights into its function at postsynaptic NMJs.

## Results

### Dyn2 forms a belt-shaped structure around the actin core of podosomes

To explore the function of Dyn2 at synaptic podosomes, first we used immunostaining and z-sectioning confocal microscopy to determine the spatial distribution of endogenous Dyn2 in myotubes differentiated from C2C12 myoblasts on laminin-coated coverslips (Kummer, Misgeld et al., 2004, Proszynski et al., 2009). Podosomes are actin-based protrusive structures comprising an actin core, an actin-based cable network, and an integrin-based adhesive ring (Fig 1a). The actin core is enriched with a branched actin polymerization machinery, with the cable network emanating from the actin core to link with the adhesive ring composed of integrin and adaptor proteins (Linder & Aepfelbacher, 2003, Linder & Wiesner, 2016, Luxenburg et al., 2007).

**Fig. 1:**
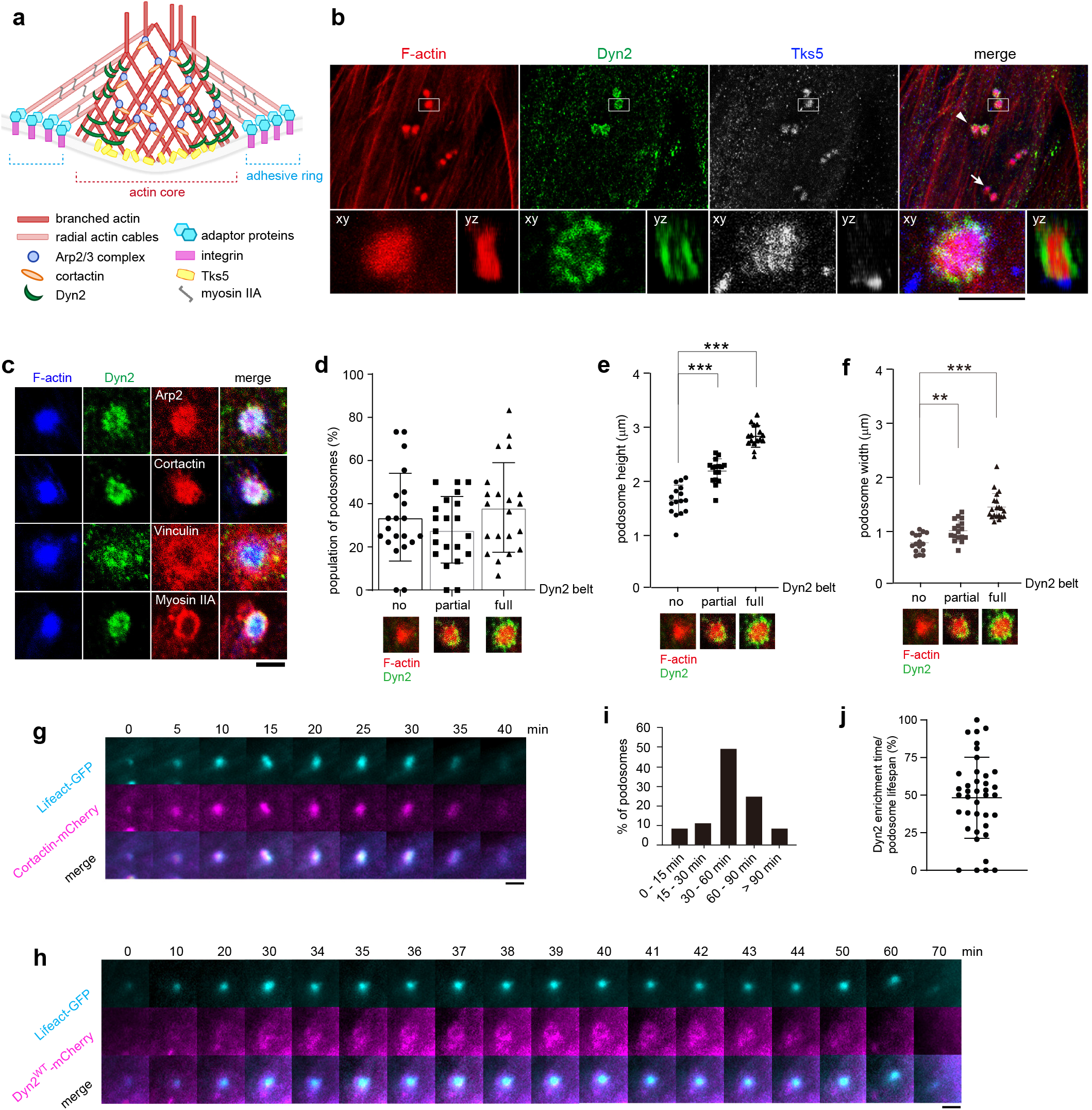
Dyn2 forms a belt-shaped structure at the actin core of podosomes. a, Schematic diagram of podosome structure. b, Dyn2 forms belt-like structures around the actin cores of synaptic podosomes. Myoblasts were seeded on laminin-coated coverslips and subjected to differentiation. Differentiated myotubes were fixed and stained to visualize endogenous Factin, Dyn2, and Tks5. Images were acquired from z-stack confocal microscopy. Boxed areas were magnified and shown in lower panels to display the Z-projection and orthogonal view of a single podosome. Arrowhead, podosome with a Dyn2 belt. Arrow, podosome without a Dyn2 belt. Scale bar, 2 μm. c, Dyn2 localizes differently from other podosome components. Differentiated myotubes were fixed and stained to visualize endogenous F-actin, Dyn2, Arp2, cortactin, vinculin, and myosin IIA. Scale bar, 2 μm. d-f, Level of Dyn2-belt formation is correlated with podosome height and width. Podosomes were grouped into three categories according to amounts of Dyn2 surrounding the actin core. Data are presented as mean ± s.d. of podosome height and width. Each dot represents one podosome. At least 40 podosomes in 10 different cells were analyzed. Statistical analyses were performed by one-way ANOVA with Dunnett’s multiple comparisons test. \*\**P* < 0.01; \*\*\**P* < 0.001. The representative images are shown in the lower panels, with Dyn2 in green and F-actin in red. g-h, Snapshots of time-lapse images of a single podosome showing the temporal distribution of Cortactin and Dyn2 in myotubes. Myoblasts were transfected with Lifeact-GFP and Cortactin-mCherry (g) or Dyn2^WT^-mCherry (h), respectively, and were seeded on laminin-coated glassbottom dishes, subjected to differentiation for 5-7 days, and imaged by inverted fluorescence microscopy at 37 °C with 1 min frame intervals. Scale bar, 2 μm. i, Frequency distribution of podosome lifespan in myotubes (n = 37 podosomes). j, Percentage of Dyn2 appearance during podosome lifespans in myotubes. Each dot represents one podosome.

Similar to a previous finding (Proszynski et al., 2009), we observed that Dyn2 is enriched at the actin core by co-staining with phalloidin and the specific podosome scaffold protein, Tks5 (Seals, Azucena et al., 2005) (Fig. 1b). An orthogonal view of a podosome from a reconstructed z-stack image further showed that Dyn2 is specifically enriched at the edge of the actin core where it forms a belt-shaped structure (yz view in Fig. 1b). In contrast, the actin polymerization machinery, comprising Arp2/3 and cortactin, was evenly enriched throughout the actin core and it partially colocalized with the Dyn2-enriched belt. Both the actin cable network and adhesive ring, labeled by myosin IIA and vinculin respectively, were located outside of the Dyn2 belt (Fig. 1c). Localization of the belt-shaped Dyn2 structure between the actin core and actin cable was further confirmed by stimulated emission depletion (STED) microscopy (Supplementary Fig. S1a, b). Therefore, despite direct interaction between Dyn2 and cortactin (McNiven, Kim et al., 2000), their slightly different localizations in podosomes suggest that Dyn2 may play an additional role in those structures apart from promoting actin polymerization. Notably, we also observed belt-shaped Dyn2 structures around actin cores of podosome rosettes in cSrc-transformed NIH3T3 fibroblasts (Supplementary Fig. S1c).

Given that Dyn2 belts were not observed in all podosomes (arrows in Fig. 1b; Supplementary Fig. S1d), we sought to investigate the role of Dyn2 in podosomes. We quantified the sizes (height and width) of actin cores and categorized podosomes into three groups—those equipped with full, partial, or no Dyn2 belt—and found that the proportion of podosomes in each group was relatively similar (Fig. 1d). Statistical analysis showed that podosomes with a Dyn2 belt had taller and wider actin cores (2.83 ± 0.20 μm in height and 1.46 ± 0.25 μm in width) compared to those with partial Dyn2 belts (2.19 ± 0.22 μm in height and 1.02 ± 0.19 μm in width) or lacking a Dyn2 belt (1.65 ± 0.28 μm in height and 0.79 ± 0.17 μm in width) (Fig. 1e, f). These findings raise the possibility that Dyn2 may contribute to podosome growth.

Next, we performed live-cell imaging to explore the temporal distribution of Dyn2 together with other critical podosome components in myotubes (Fig. 1g, h). Consistent with the kinetics of podosome components in other cell types (Luxenburg, Winograd-Katz et al., 2012), we found that F-actin and cortactin accumulated synchronously during initiation of synaptic podosomes, which was followed by Tks5 recruitment (Fig. 1g; Supplementary Fig. S1e). This result supports the roles of cortactin in podosome initiation/formation and Tks5 in podosome maturation. Defining podosome lifespan according to the period of F-actin appearance, we found that podosomes in myotubes exhibited heterogenous lifespans, with >90% of podosomes having lifetimes longer than 15 min, which differs from the short-lived (<15 min) podosomes in macrophages and osteoclasts (Destaing, Saltel et al., 2002, Guiet, Verollet et al., 2012) (Fig 1i, Supplementary video 1). Furthermore, more than 30% of the podosomes we assessed had a lifespan longer than 60 min. Dyn2 enrichment possesses ~48 % of the lifespan of podosomes (Fig. 1j; Supplementary Fig. S1f), and transient Dyn2 appearance explained the occurrence of podosomes decorated with partial Dyn2 belts at a given time-point. We also noticed that, after a Dyn2 belt disappeared, the actin core (labeled by Lifeact-GFP) gradually dissembled and eventually vanished, suggesting a potential role for Dyn2 in podosome turnover.

### Dyn2 is required for the growth of podosomes

To investigate the function of Dyn2 in podosomes, we downregulated Dyn2 in C2C12-derived myotubes by means of lentiviral shRNAs (Fig. 2a; Supplementary Fig. S2a) and then examined the morphology of podosomes by z-stack confocal microscopy. Dyn2 knockdown significantly reduced both the mean height and width of podosome cores from 1.81 μm and 1.52 μm to 1.03 μm and 0.86 μm, respectively, without affecting podosome density (Fig. 2b, c). Consistent with the defect in podosome growth, podosome lifespan was also significantly diminished in Dyn2-depleted myotubes (Fig. 2d, e). These results demonstrate an essential role for Dyn2 in podosome growth, but not podosome initiation.

**Fig. 2:**
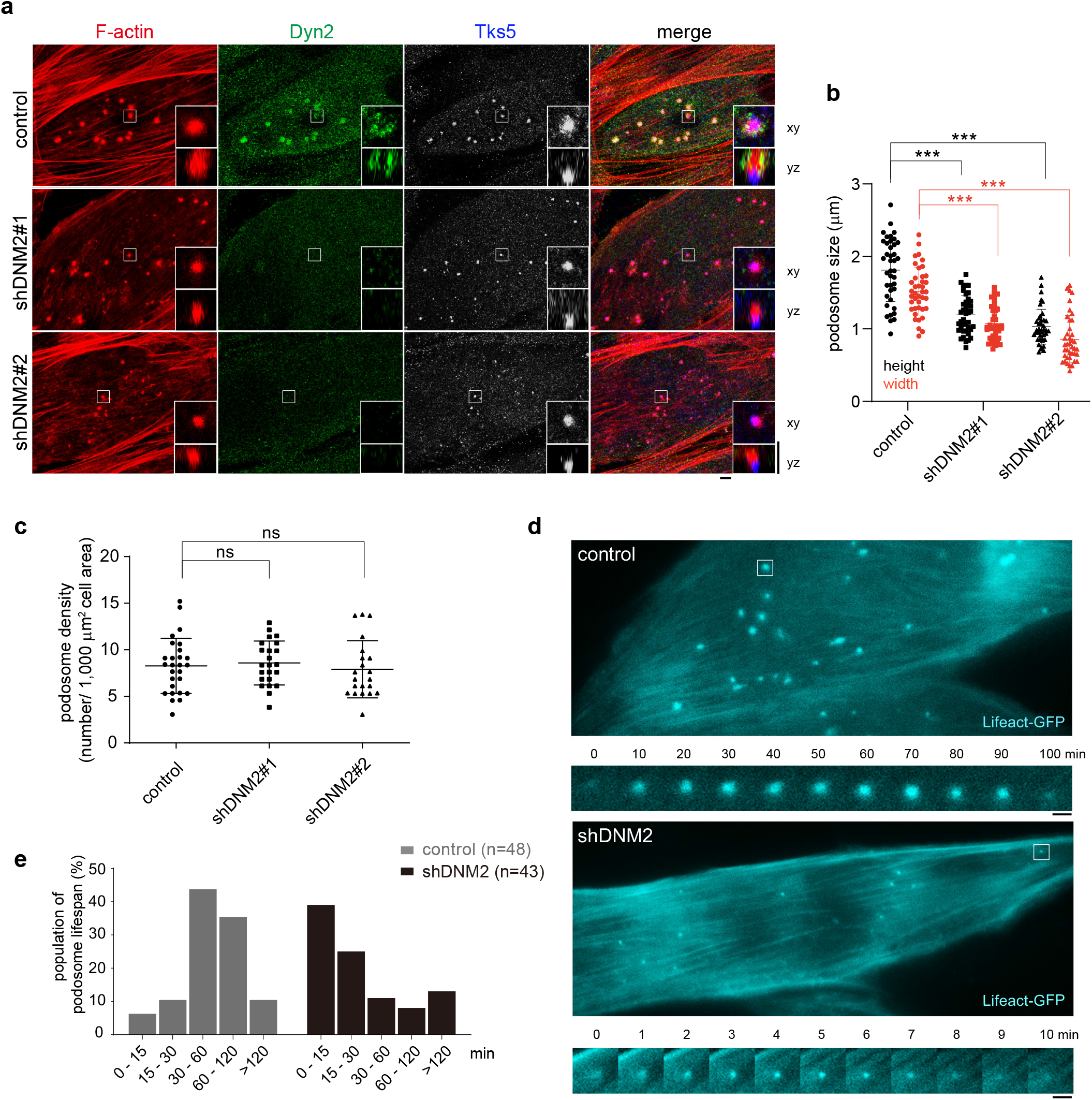
Dyn2 is required for podosome growth. a-c, Podosomes in Dyn2-depleted myotubes are smaller than in control cells. Podosomes were labeled with the podosome marker, Tks5. Day 5-differentiated myotubes were fixed and stained to visualize endogenous F-actin, Dyn2, and Tks5. Images were acquired by z-stack confocal microscopy. Insets represent enlarged views of Z-projection (upper) and orthogonal views (lower) of a single podosome. Scale bars, 2 μm. b, Height and width of podosome actin cores are reduced in Dyn2-depleted myotubes. Each dot represents one podosome. c, Podosome density in Dyn2-depleted myotubes. Each dot represents one myotube. Data are presented as mean ± s.d. de, Podosomes in Dyn2-depleted myotubes have shorter lifespans. Podosome lifespan was quantified by the duration of the podosome core, as labeled by Lifeact-GFP, and the data was divided into five cohorts as shown.

In contrast, acute treatment with the dynamin GTPase inhibitor dynasore induced an accumulation of Dyn2 at podosomes, which resulted in time-dependent actin core elongation (Supplementary Fig. S2b, c), suggesting that the GTPase activity of Dyn2 is critical for podosome turnover. However, we noted that prolonged dynasore incubation (12 h) resulted in a reduction of podosome numbers in myotubes, likely due to indirect effects or a pleiotropic cellular response caused by blockage of dynamin-mediated endocytic pathways (Supplementary Fig. S2d).

### Dysregulated Dyn2 activity affects podosome growth and turnover

To understand which biochemical activity of Dyn2 is responsible for the growth of podosomes, we utilized adenovirus to transiently overexpress different dominant-negative mutants of Dyn2 in wild-type or Dyn2-depleted myotubes. These mutants included a hyper-assembling CNM-linked mutant (A618T), a membrane fission-defective CMT-linked mutant (G537C), a mutant with lower actin binding ability (K/E) or with higher actin binding activity (E/K), as well as a GTPase-defective mutant (K44A) (Chin et al., 2015, Gu et al., 2010, Kenniston & Lemmon, 2010) (Fig. 3a, b). Further detailed information and references for these mutants are provided in Fig. 3b and Supplementary Table 1.

**Fig. 3:**
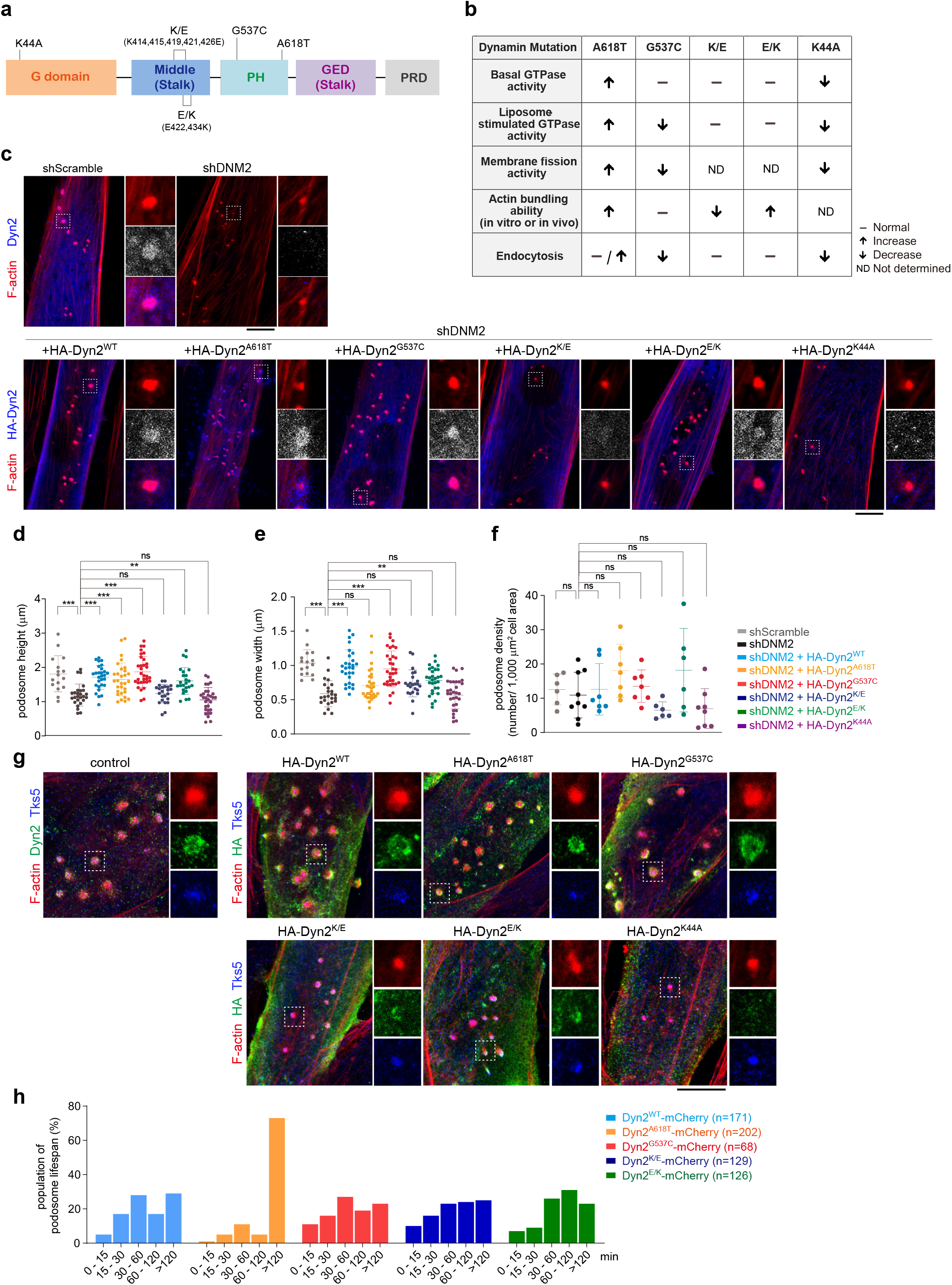
Dyn2 is essential for podosome growth and turnover. a, Domain structure of Dyn2 and the mutations used in this experiment. b, Dyn2 mutants and their biochemical properties. c, Effects of different HA-tagged Dyn2 mutants on rescuing the morphology of podosomes in Dyn2-knockdown myotubes. Endogenous Dyn2 was stained in control cells. HA-tagged Dyn2, F-actin, and Tks5 were stained and imaged under z-stack confocal microscopy after 16 h of induction. d-e, Height and width of podosomes. Each dot represents one podosome (n = 16~32 podosomes per mutant). For each mutant, 6~9 different cells were analyzed. f, Podosome density in myotubes. Each dot represents the podosome density in one cell. Statistical analyses were performed by one-way ANOVA with Dunnett’s multiple comparisons test. ns, not significant; \*\**P* < 0.01; \*\*\**P* < 0.001. g, Subcellular localization of different HA-tagged Dyn2 mutants in wild-type myotubes. h, Frequency distribution of podosome lifetime in myotubes expressing different Dyn2 mutants. Myoblasts were transfected with Lifeact-GFP and Dyn2-mCherry to visualize F-actin and Dyn2, respectively. Myoblasts were seeded on laminin-coated glass-bottom dishes, subjected to differentiation for 5-7 days, and imaged by inverted fluorescence microscopy at 37 °C with 2 min frame intervals. Lifespans of individual podosomes were tracked and measured manually. For each condition, at least 68 podosomes for each mutant were analyzed.

Exogenously-expressed wild type Dyn2 (Dyn2^WT^), Dyn2^G537C^ and Dyn2^E/K^ were enriched in podosomes and rescued the size of podosome actin cores in Dyn2-depleted myotubes (Fig. 3c-f). In contrast, neither Dyn2^K/E^ nor Dyn2^K44A^ were enriched in podosomes, nor could they rescue the size of actin cores. Intriguingly, Dyn2^A618T^ did localize to podosomes and restored actin core height but not width. Similar effects of these Dyn2 mutants on synaptic podosomes were also observed in wild type myotubes, with Dyn2^A618T^, Dyn2^G537C^ and Dyn2^E/K^ localizing to podosomes, but Dyn2^K/E^ and Dyn2^K44A^ not doing so (Fig. 3g; Supplementary Fig. S3a-d). Importantly, the Dyn2^A618T^ mutant protein (which exhibits hyper-self-assembly) led to a significant increase in podosome height, whereas the membrane fission-defective Dyn2^G537C^ mutant protein did not affect any apparent morphological feature of podosomes. These results suggest that the actin binding activity of Dyn2 is critical for its enrichment at podosomes, and hyper-self-assembly of Dyn2 can result in abnormal podosome morphology.

To further examine the effect of Dyn2 on podosome turnover, we co-transfected LifeAct-GFP and mCherry-tagged Dyn2 mutants into myoblasts and induced their differentiation into myotubes to measure the lifespan of podosomes. Statistical analysis revealed that whereas expression of Dyn2^G537C^-mCherry, Dyn2^K/E^-mCherry, and Dyn2^E/K^-mCherry did not significantly alter podosome lifespan, Dyn2^A618T^-mCherry expression prolonged podosome lifetimes, manifesting as a dramatic increase in long-lived podosomes (> 2 hr in Fig. 3h). Moreover, unlike the temporal emergence of Dyn2^WT^ in podosomes (Fig. 1h), Dyn2^A618T^ belts were very stable throughout the lifetime of podosomes (Supplementary video 2). In Supplementary Fig. S3e, we present an example of podosomes decorated with Dyn2^A618T^ belts for 14.5 hr with an overall lifetime of ~15 hr.

Regrettably, we were unable to analyze the effect of Dyn2^K44A^-mCherry on podosome lifetime in myotubes due to its strong dominant-negative effect on endocytosis and myoblast differentiation (Chuang, Lin et al., 2019), which prevented us from generating myotubes that expressed Dyn2^K44A^-mCherry. Instead, we analyzed the effect of Dyn2 mutants on podosome rosettes in cSrc-transformed NIH3T3 fibroblasts and observed an increase in podosome height and lifespan in Dyn2^A618T^-expressing cells, whereas there was a notable decrease in podosome size and lifespan in Dyn2^K/E^- and Dyn2^K44A^-expressing cells (Supplementary Fig. S3f-i). Together, these results show that the actin binding, self-assembly, and GTP hydrolysis activities of Dyn2, but not its membrane fission ability, are involved in regulating podosome growth and turnover.

### Dyn2 is required for synaptic podosome function

The function of synaptic podosomes is to promote NMJ development through their ECM degradative ability, a critical feature of mature podosomes (Chan et al., 2020, Linder, 2007, Proszynski et al., 2009). To assess if proper activity of Dyn2 is required for the functionality of synaptic podosomes, we performed an ECM degradation assay on Dyn2-depleted myotubes re-expressing either wild-type or different mutant proteins. Similar to their effects on podosome morphology, Dyn2^WT^, Dyn2^A618T^, Dyn2^G537C^ and Dyn2^E/K^ could restore the ECM degradative ability of Dyn2 knockdown myotubes, but Dyn2^K/E^ or Dyn2^K44A^ mutants could not (Fig. 4a, b). Furthermore, expression of Dyn2^A618T^ in wild type myotubes resulted in a significant increase in the area of degradation (Fig. 4c, Supplementary Fig. S4a). Given that podosome density in Dyn2^A618T^-expressing myotubes remained unchanged (Supplementary Fig. S3d), this increase in ECM degradation indicates that those podosomes enclosed by Dyn2^A618T^ not only have taller actin cores and longer lifetimes, but also exhibit a better ECM degradative ability. Thus, our results demonstrate that Dyn2 promotes podosome maturation and turnover, and that the CNM-associated Dyn2^A618T^ mutant protein has a dominant effect on the stability of synaptic podosomes.

**Fig. 4:**
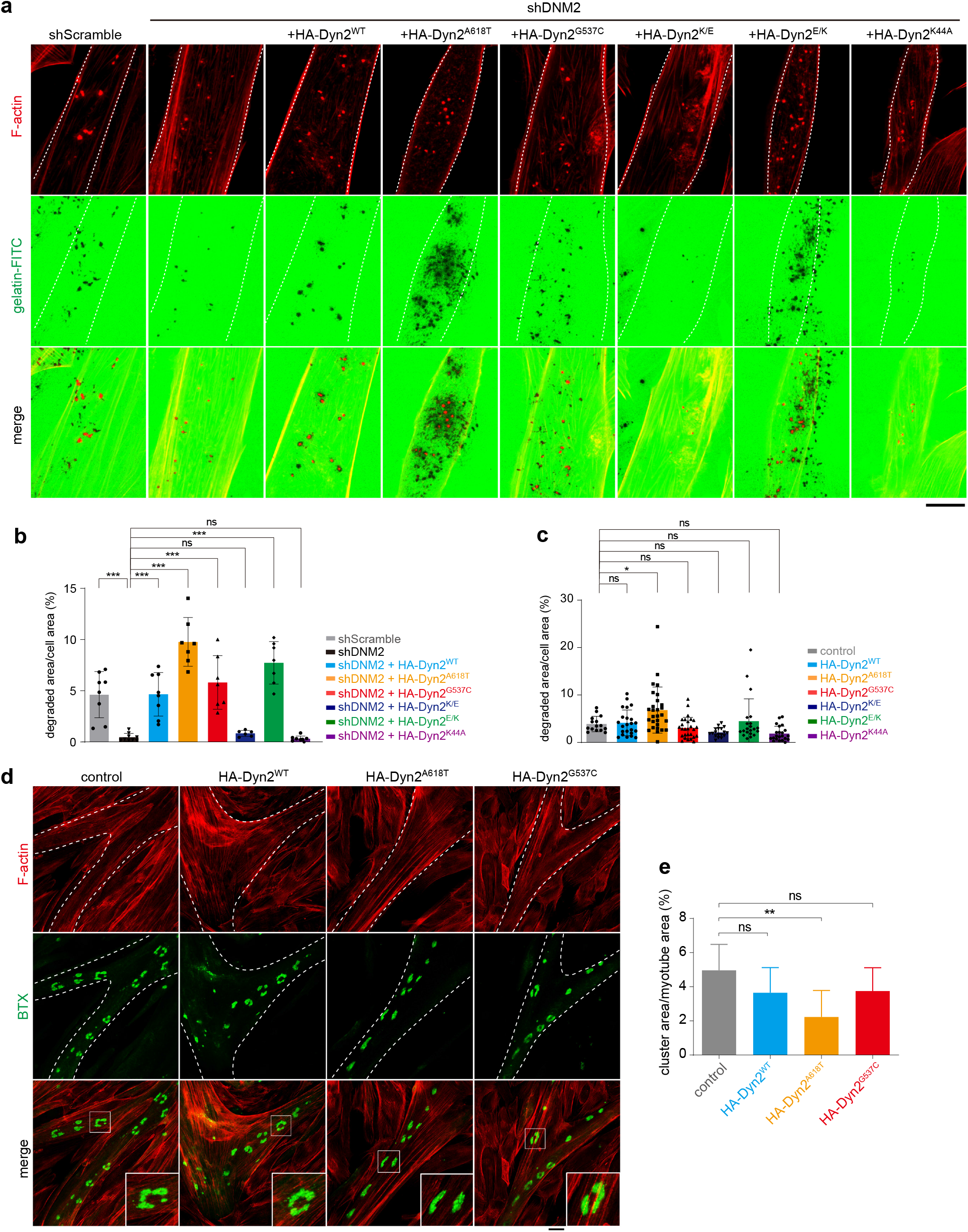
Dyn2 regulates the matrix degradation activity of muscle podosomes and postsynaptic AChR clusters. a, Rescue effects of HA-tagged Dyn2 mutants on the ECM degradation activities of Dyn2-depleted myotubes. HA-tagged Dyn2 and F-actin in Day 4 Dyn2-KD myotubes were stained and imaged under confocal microscopy after 16 h of induction and culturing on gelatin-FITC-coated coverslips. Dashed lines indicate cell boundaries. Scale bar, 20 μm. b, Quantification of degraded area versus cell area, shown as mean ± s.d. Each dot represents the percentage of degraded area in each cell. For each mutant, at least six different cells were analyzed. c, Quantification of gelatin-FITC degraded area of HA-tagged Dyn2 mutants expressing wild-type myotubes. Each dot represents the percentage of degraded area in each cell. For each mutant, at least 15 different cells were analyzed. d. HA-tagged Dyn2, F-actin, and AChR (labeled by Bungarotoxin) in Day 6 myotubes cultured on Permanox slides in eight-well Flexiperm chambers, stained and imaged under z-stack confocal microscopy after 16 h of induction. Dashed lines indicate cells expressing HA-tagged Dyn2 mutants. Scale bar, 20 μm. e, Quantification of total AChR area versus cell area, shown as mean ± s.d. For each mutant, at least eight different cells were analyzed. Statistical analyses were performed by one-way ANOVA with Dunnett’s multiple comparisons test. \**P* < 0.05, \*\**P* < 0.01.

To further explore the effect of Dyn2 on synaptic podosome-mediated NMJ development, we assessed AChR distribution by staining with Alexa488-conjugated bungarotoxin (BTX). The AChR clusters in Dyn2^WT^ and Dyn2^G537C^ myotubes were perforated and had similar areas relative to control, whereas the AChR clusters in Dyn2^A618T^-expressing myotubes were perforated but with less cluster area (Fig. 4d, e). Decreased AChR area in Dyn2^A618T^-expressing myotubes might be due to higher ECM remodeling ability of Dyn2^A618T^ (Fig. 4c). This finding suggests that Dyn2 plays a critical role in synaptic podosome and AChR cluster maturation, and that the CNM-associated Dyn2 mutant variant may perturb their development.

### Dyn2 regulates actin-based postsynaptic cytoskeletal organization and postsynaptic development

To establish if Dyn2^A618T^ could affect the development and electrophysiological function of NMJs, we utilized *Drosophila* as it is a commonly used system for studying synaptic development and function. We generated *UAS* transgenes of HA-tagged wild-type and mutant human Dyn2, and expressed each of them predominantly in muscles using the *MHC-GAL4* driver (Schuster, Davis et al., 1996). We observed that Dyn2^WT^ was localized in postsynaptic NMJs and colocalized with α-spectrin, an actin-binding protein associated with the postsynaptic plasma membrane and that is involved in establishing the postsynaptic subsynaptic reticulum (SSR) (Supplementary Fig. S5a) (Pielage, Fetter et al., 2006). The Dyn2^A618T^ and Dyn2^G537C^ mutant variants also presented similar enrichment at postsynaptic NMJs, but the pattern and intensity of α-spectrin signal was significantly reduced upon expression of Dyn2^A618T^ (Fig. 5a-b, Supplementary Fig. S5a), even though expression levels of α-spectrin remained constant among all these flies (Supplementary Fig. S5b). Unlike the disrupted pattern of spectrin signal in Dyn2^A618T^-expressing NMJs, other NMJ components were not significantly affected, such as the SSR marker DLG and the active zone marker Brp (Supplementary Fig. S5c, d), indicating that Dyn2^A618T^ specifically affects the organization of the postsynaptic actin cytoskeleton.

**Fig. 5:**
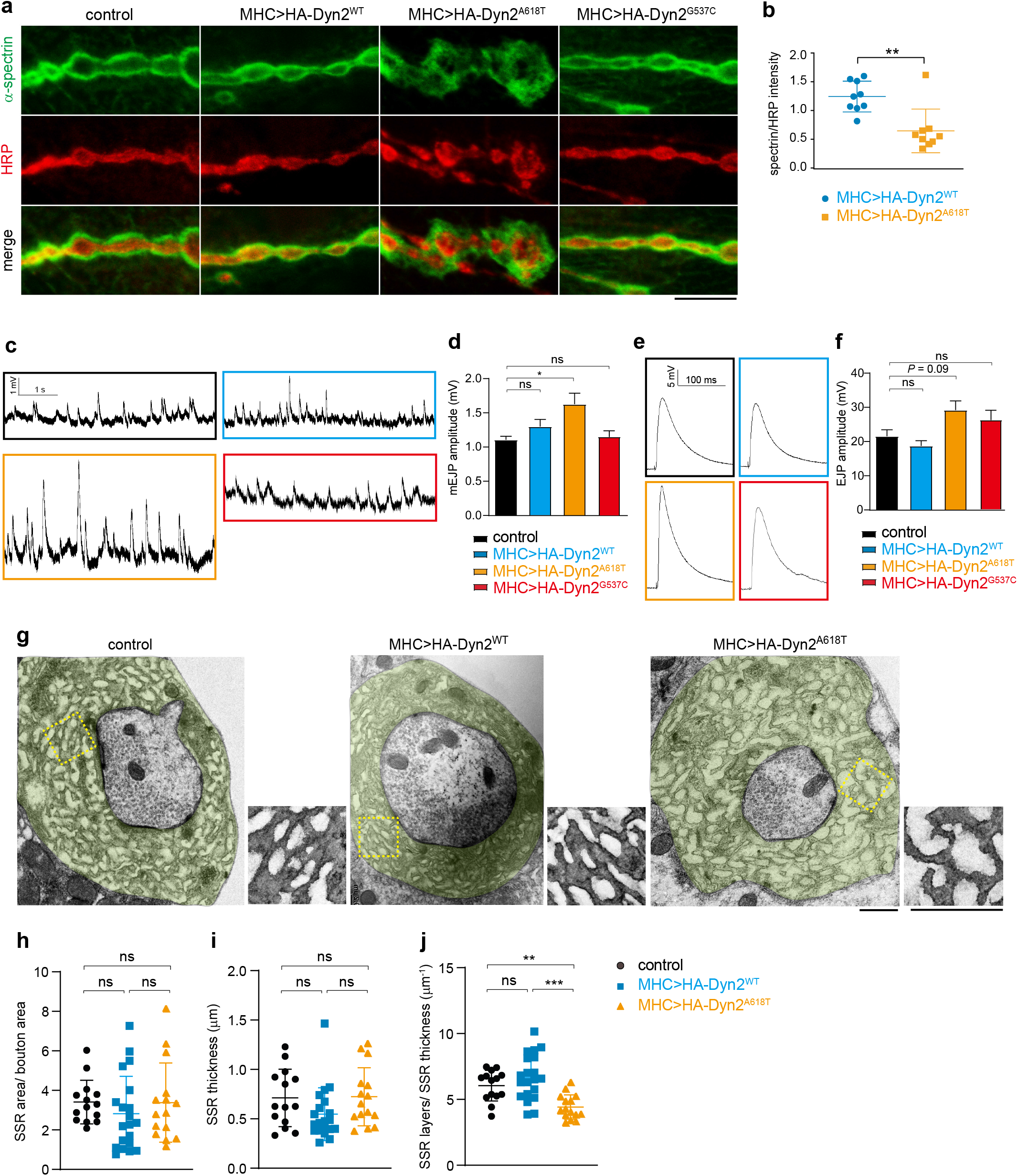
Dyn2 is involved in the development and activity of *Drosophila* NMJs. a, Distribution of α-spectrin and HRP in third-instar larval NMJs, as revealed by immunostaining with anti-α-spectrin and HRP antibodies. Images were taken by z-stack confocal microscopy and are presented as z-stack projections. Scale bar, 10 μm. b, The intensity of α-spectrin normalized to HRP intensity. Each dot represents the normalized intensity in one NMJ. Statistical analysis was performed by Student’s t-test. \*\**P* < 0.01. For each mutant, nine NMJs were analyzed. c-f, Representative traces and quantification of the mEJP amplitude and EJP amplitude of wild type and *Drosophila* expressing Dyn2^WT^ and Dyn2^A618T^ in their body wall muscle. g, TEM images of NMJ boutons from wild type and Dyn2^WT^- and Dyn2^A618T^-expressing *Drosophila*. Boxed areas (within yellow dashed lines) are magnified and are shown alongside. Scale bar, 500 nm. The SSR is pseudocolored in green, and parameters of SSR morphology were quantified and are shown in h-j. Each dot represents one bouton. Statistical analysis of panel d,f and h-j was performed by one-way ANOVA with Tukey’s multiple comparisons test. \**P* < 0.05, \*\**P* < 0.01, \*\*\**P* < 0.001.

During the development of *Drosophila* NMJs, α-and β-spectrins are associated with the postsynaptic plasma membrane, where they form a complex with F-actin for SSR development, with RNAi knockdown of α- or β-spectrin altering SSR membrane organization and synaptic transmission (Pielage et al., 2006). To examine the functional consequences of altered spectrin organization, we recorded the membrane potential of postsynaptic NMJs from Dyn2^A618T^-expressing *Drosophila* larvae and observed increased amplitudes of miniature excitatory junction potential (mEJP) and evoked EJP (Fig. 5c-f), reminiscent of the effect of spectrin depletion in *Drosophila* (Pielage et al., 2006). However, mEJP frequency and Quantal content were not altered in these mutants (Supplementary Fig. S5e, f).

Greater mEJP amplitude could result from an increased level or activity of postsynaptic glutamate receptors, or both. However, we noted that neither the expression level nor the cluster size of GluRIIA, the major subunit of glutamate receptors (Marrus, Portman et al., 2004), was altered by expression of Dyn2^A618T^ (Supplementary Fig. S5g-j). Given that expression of Dyn2^A618T^ alters postsynaptic organization and electrophysiological activities, we examined SSR ultrastructure using transmission electron microscopy (TEM) and observed that Dyn2^WT^ or Dyn2^A618T^ did not cause any apparent change in the area or thickness of SSR (Fig. 5f-h). However, expression of Dyn2^A618T^, but not Dyn2^WT^, reduced SSR density (SSR layers/SSR thickness), indicative of a looser membrane structure (Fig. 5f, i). Thus, our ultrastructural and electrophysiological data indicate that the hyper self-assembly activity of the Dyn2^A618T^ mutant protein alters actin-dependent cytoskeletal organization, thereby disturbing the electrophysiological function of the postsynaptic compartment of *Drosophila* NMJs. It is worth noting that, similar to *Drosophila* NMJs, endogenous Dyn2 is also enriched at mouse NMJs (Supplementary Fig. S5k, l).

### The actin bundling activity of Dyn2 is regulated by GTP hydrolysis

To directly investigate how Dyn2 remodels actin cytoskeleton, we utilized *in vitro* reconstitution to explore the aforementioned biochemical activities of Dyn2 and its effect on actin bundle formation. Similar to previous reports (Chuang et al., 2019, Gu et al., 2010), we observed prominent F-actin bundling activity for Dyn2, but much less bundled actin for Dyn1 (Fig. 6a, b; Supplementary Fig. S6a, b). Strikingly, Dyn2-actin bundles were significantly reduced upon addition of GTP, but not the non-hydrolyzable GTP analog, GMPPCP (Fig 6 c, d). We visualized Dyn2-mediated actin bundling and the disassembly induced by GTP in real-time by imaging rhodamine-labeled actin under fluorescence microscopy (Fig. 6e).

**Fig. 6:**
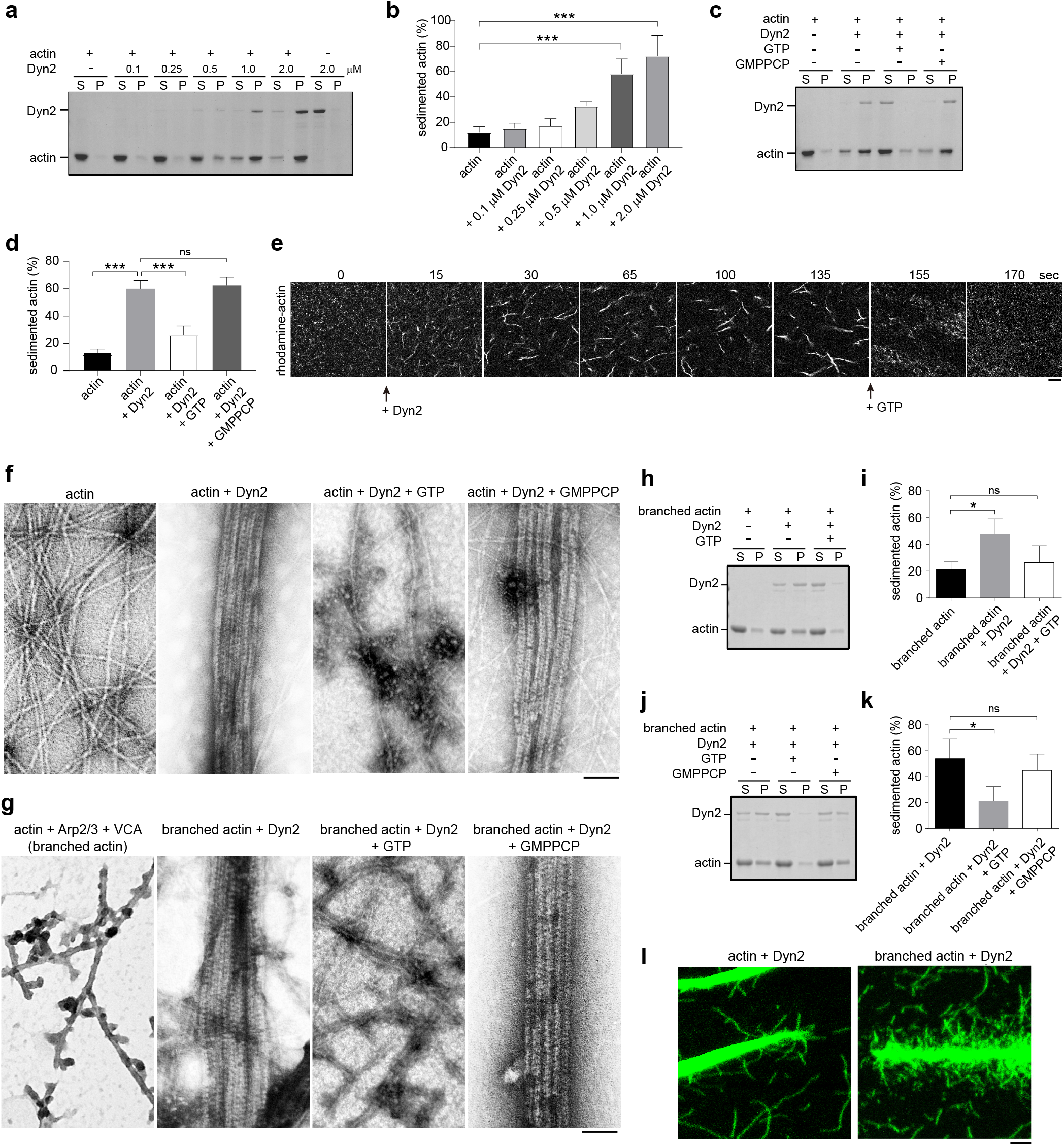
Dyn2 bundles both linear and branched actin filaments. a, Dyn2 bundles actin filaments. Reconstituted F-actin (5 μM) was incubated with Dyn2 at indicated concentrations for 30 min and subjected to 20 min centrifugation at 14,000 x *g*. S, supernatant; P, pellet. b, Dyn2 (1 μM) shows significant actin bundling activity. Mean ± s.d. of percentage of sedimented actin was quantified in ImageJ as the ratio of actin in the pellet versus total actin. c-d, F-actin sedimentation assay and quantification result of Dyn2 bundling activity. Reconstituted F-actin (5 μM) was incubated with 1 μM Dyn2 for 30 min followed by 15 min incubation with 1 mM GTP or GMPPCP. e, Real-time visualization of the actin bundling activity of Dyn2 under confocal microscopy. Scale bar, 10 μm. f, TEM images of negative-stained actin-Dyn2 bundles. Scale bar, 100 nm. g, TEM images of negative-stained branched actin-Dyn2 bundles. Scale bar, 100 nm. h-k, Branched F-actin sedimentation assay and quantification result of Dyn2 bundling activity. Concentrations of individual components were 5 μM actin, 1 μM Dyn2, 30 nM Arp2/3 complex, 80 nM VCA and 1 mM nucleotides. Statistical analyses were performed by one-way ANOVA with Dunnett’s multiple comparisons test. ns, not significant; \**P* < 0.05; \*\*\**P* < 0.001. l, Morphology of Dyn2-bundled linear and branched actin under confocal microscopy. Scale bar, 1 μm.

To better visualize how addition of GTP affects Dyn2-mediated actin bundling, we utilized negative-stain TEM to image the Dyn2-actin bundles with or without nucleotides. We found that unlike the aligned and bundled actin filaments surrounded by ordered Dyn2 rings in the absence of GTP, the Dyn2 rings were disassembled and the actin bundles became dispersed upon addition of GTP, but not GMPPCP (Fig. 6f). We also analyzed the effect of other GTP analogs, including GDP and GDP:AlF_4_^-^ (which mimics the transition state of GTP hydrolysis), and found that actin bundles remained stable, but Dyn2-actin assemblages were less ordered in the presence of GDP relative to GMPPCP (Supplementary Fig. S6c).

The GTP hydrolysis-induced Dyn2 disassociation from actin is reminiscent of Dyn2 disassembly from the membrane upon GTP hydrolysis (Bashkirov, Akimov et al., 2008, Chin et al., 2015, Pucadyil & Schmid, 2008). Furthermore, we occasionally observed a partially packed Dyn2-actin bundle under TEM, perhaps representing an intermediate stage of the bundling process (Supplementary Fig. S6e). Images from 3D tomography clearly show that Dyn2 assembles around actin filaments to form a packed actin bundle (Supplementary Fig. S6f and video 3). Note, we sometimes observed Dyn2 rings inside the actin bundle, indicating that *in vitro* reconstitution assays should be conducted carefully and validated with *in vivo* experiments.

Since the actin core of podosomes is predominantly composed of branched actin (Albiges-Rizo, Destaing et al., 2009), we repeated our actin sedimentation assays with branched actin reconstituted by Arp2/3 complex and the VCA domain of neural Wiskott-Aldrich syndrome protein (N-WASP) to examine the bundling ability of Dyn2 on branched actin. We found that Dyn2 displays prominent bundling activity on branched actin, which is also regulated by GTP hydrolysis (Fig. 6g-k). The morphological differences of Dyn2-bundled linear and branched actin were better visualized under confocal microscopy (Fig. 6l). Together, these results show that Dyn2 bundles both types of actin filaments that are present in podosomes, and that assembly of Dyn2 oligomers on actin filaments is regulated by GTP hydrolysis.

### CNM-associated Dyn2 mutants are insensitive to GTP hydrolysis

Given the differential effect of Dyn2^A618T^ and Dyn2^G537C^ on cytoskeleton remodeling *in vivo* (Fig. 5a), we examined if these two mutants exert different impacts on actin bundles. Using an actin sedimentation assay, we found that both Dyn2^A618T^ and Dyn2^G537C^ present comparable actin bundling activities (Fig. 7a, b). However, whereas actin filaments bundled by Dyn2^WT^ and Dyn2^G537C^ dissociated upon GTP hydrolysis, those bundled by Dyn2^A618T^ were resistant to GTP addition and remained bundled (Fig. 7a, b). Importantly, both linear and branched actin bundled by other CNM-associated Dyn2 mutant proteins, Dyn2^R465W^ and Dyn2^S619L^, were also insensitive to GTP (Fig. 7c-g, Supplementary Fig. S7). These findings indicate that Dyn2 can bundle both unbranched and branched actin, which is terminated by GTP hydrolysis. Thus, Dyn2 assembly and sensitivity to GTP hydrolysis play decisive roles in actin organization, and these biochemical features are critical for podosome turnover as well as NMJ morphology.

**Fig. 7:**
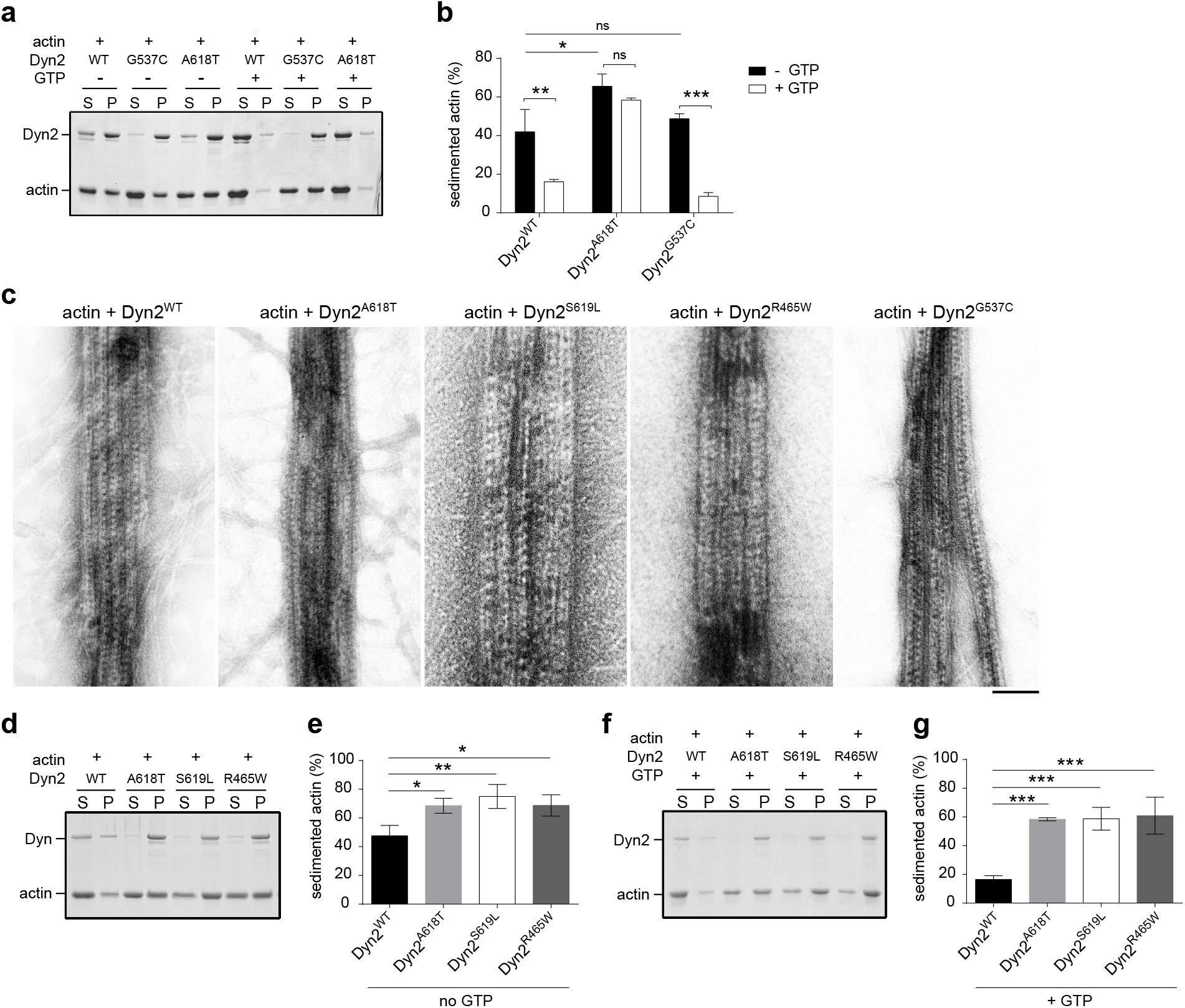
Actin filaments bundled by hyper self-assembled Dyn2 mutants are less sensitive to GTP. a, Actin bundles assembled by Dyn2^A618T^ are resistant to GTP hydrolysis. Reconstituted F-actin (5 μM) was incubated with 1 μM Dyn2 for 30 min, followed by 15 min incubation with 1 mM GTP, and subjected to low-speed centrifugation at 14,000 x *g*. S, supernatant; P, pellet. b, Mean ± s.d. of the percentage of sedimented actin was quantified in ImageJ as the ratio of actin in the pellet versus total actin. c, TEM images of negative-stained actin-Dyn2 bundles. Scale bar, 100 nm. d-g, F-actin sedimentation assay and quantification of Dyn2 bundling activity in the absence of GTP (d,e) or in the presence of GTP (f,g). Statistical analyses were performed by one-way ANOVA with Dunnett’s multiple comparisons test. ns, not significant; \**P* < 0.05; \*\**P* < 0.01; \*\*\**P* < 0.001.

### Phosphorylation of Dyn2 residue Y597 is important for its podosome targeting and actin bundling activity

Dyn2 is recruited to plasma membrane clathrin-coated pits by interacting with several SH3 domain-containing proteins (Meinecke, Boucrot et al., 2013). However, despite there being a direct interaction between Dyn2 and the podosome components cortactin and Tks5 (McNiven et al., 2000, Oikawa, Itoh et al., 2008), we have already shown that the spatiotemporal distribution of Dyn2 is somewhat distinct from those core components (Fig. 1), raising the possibility that targeting of Dyn2 to podosomes may be regulated by a mechanism other than protein-protein interactions. Src tyrosine kinase is known to be a critical podosome-initiating enzyme (Gimona, Buccione et al., 2008, Murphy & Courtneidge, 2011). Dyn2 has been reported to be a substrate of Src kinase and is phosphorylated at residue Y597, both *in vivo* and *in vitro* (Ahn, Kim et al., 2002). Interestingly, Src-mediated phosphorylation was shown to induce Dyn1 self-assembly *in vitro*, as well as promote the function of Dyn2 on Golgi apparatus (Ahn et al., 2002, Weller, Capitani et al., 2010).

To test if Y597 phosphorylation of Dyn2 is responsible for its targeting to podosomes, we expressed a phospho-deficient mutant fusion construct, Dyn2^Y597F^-mCherry, in myotubes and observed a significant reduction in podosome association for this mutant (Fig. 8a, b). We also noted that expression of a phospho-mimetic mutant, Dyn2^Y597E^, interfered with myoblast differentiation, thus preventing us from examining its effect on synaptic podosomes. Instead, we used cSrc-transformed NIH3T3 fibroblasts to examine the effect of mutant Dyn2^Y597^. Similar to its effect in myotubes, Dyn2^Y597F^ also presented reduced enrichment at podosomes and induced smaller podosome rosettes, whereas Dyn2^Y597E^ clearly targeted to podosome rosettes (Supplementary Fig. S8a, b). Moreover, in the actin sedimentation assay, more of the bundled actin assembled by phospho-mimetic Dyn2^Y597E^ remained in the presence of GTP, indicating a slower dissociation rate of this mutant variant (Fig. 8c, d). Consistent with that outcome, much Dyn2^Y597E^-bundled actin could still be observed under TEM upon GTP treatment (Fig. 8e). Similar to Src-phosphorylated Dyn1 (Ahn et al., 2002), Dyn2^Y597E^ also displayed enhanced self-assembly in low salt buffer (Supplementary Fig. S8c), recapitulating the hyper self-assembly of the Dyn2^A618T^ mutant. Accordingly, our results demonstrate that Y597 phosphorylation is the molecular trigger for the actin bundling activity and podosome targeting of Dyn2.

**Fig. 8:**
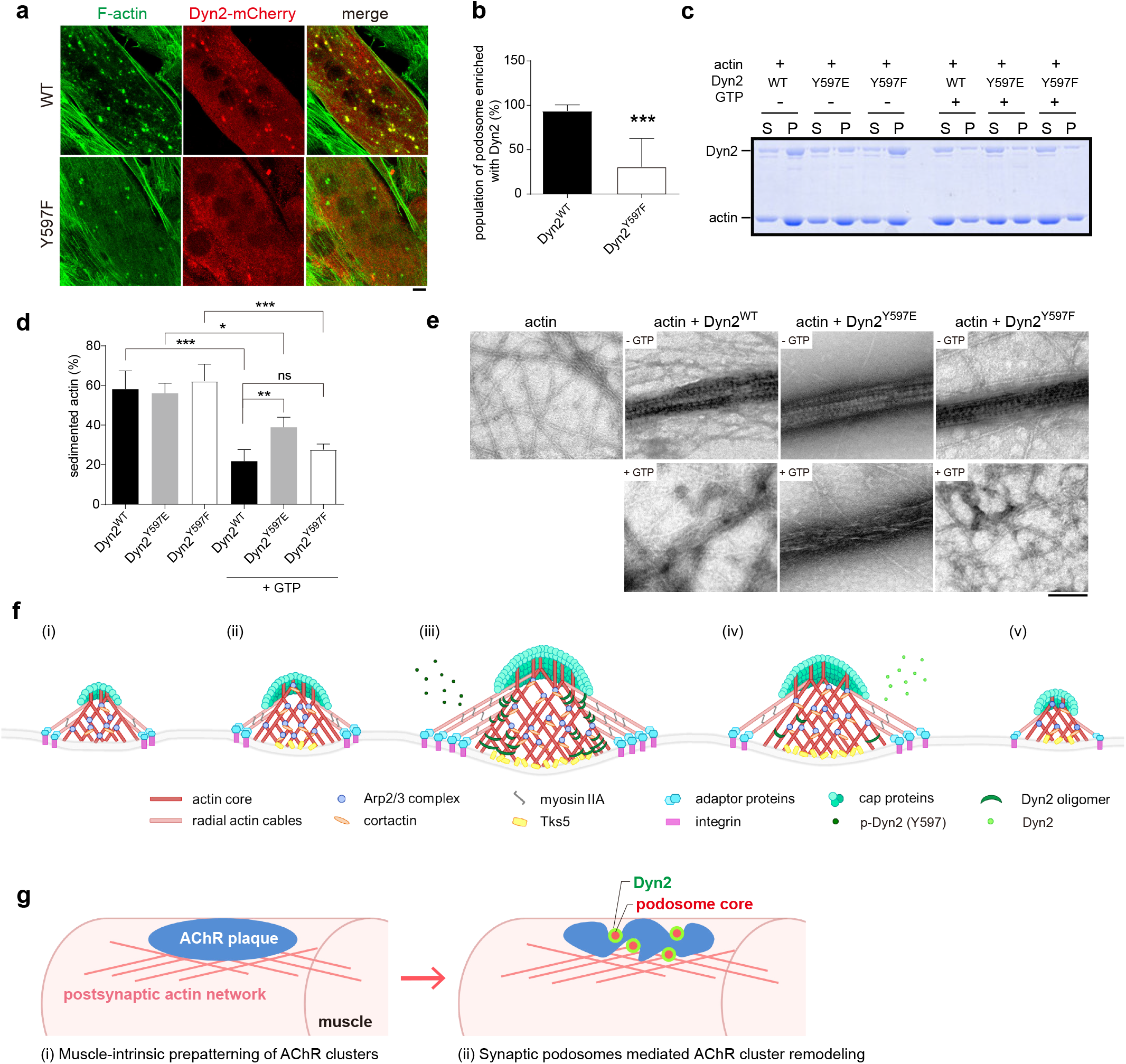
Phosphorylation of Dyn2 residue Y597 is important for its podosome targeting and actin bundling ability. a, Distribution of phospho-deficient Dyn2^Y597F^-mCherry in myotubes. b, Percentage of synaptic podosomes with Dyn2-mCherry enrichment in myotubes (n = 19 podosomes per condition). c, Actin bundling ability of Dyn2 mutants with or without GTP. The percentage of sedimented actin was quantified and is shown in d. e, TEM images of negative-stained actin-Dyn2 bundles with or without 15 min GTP incubation. Scale bar, 100 nm. f, Dyn2 functions as a molecular girdle and checkpoint for podosome maturation and turnover, respectively. (i) Interactions between ECM and integrins initiate podosome formation. Actin polymerization proteins such as Arp2/3 complex and cortactin drive podosome initiation. (ii) Tks5 is recruited to the nascent podosome and promotes podosome maturation. (iii) Phosphorylated Dyn2 is recruited to the podosome to bundle the actin core and facilitate its growth. Dyn2 forms a belt-like structure around the actin core to enhance the function of podosome. (iv, v) After dephosphorylation, GTP hydrolysis triggers Dyn2 dissociation from the podosome and induces its turnover. g, Functional role of Dyn2 in postsynaptic NMJ morphogenesis. (i) During NMJ prepatterning, AChR clusters are induced by extracellular signals, such as laminin or Wnt. (ii) During NMJ development, perforation and remodeling of AchR clusters is facilitated by synaptic podosomes whose maturation and turnover is governed by Dyn2.

## Discussion

In this study, we reveal a mechanochemical role for Dyn2 in synaptic podosome maturation and turnover, which facilitates the development of NMJs. We provide evidence of Dyn2 enrichment and function at the NMJs of *Drosophila* and mouse, as well as its role in postsynaptic membrane development during synaptogenesis. Upon phosphorylation of residue Y597, Dyn2 directly targets to and bundles actin filaments to form belt-shaped structures around podosomal actin cores, which promotes podosome maturation, whereas GTP hydrolysis-induced Dyn2 disassembly triggers podosome turnover (Fig. 8f). Thus, Dyn2 activity is involved in the development and electrophysiological activity of NMJs through its regulation of synaptic podosome maturation and turnover (Fig. 8g).

An essential role for Dyn2 in actin organization was demonstrated a decade ago (Bruzzaniti, Neff et al., 2005, Destaing, Ferguson et al., 2013, Ochoa, Slepnev et al., 2000). It was generally assumed that Dyn2 functions to promote actin polymerization through its abilities to remove capping proteins and direct interactions with cortactin, Nck, and profilin (Gu et al., 2010, Mooren et al., 2009, Schafer, 2004). Here, we discovered a structural role for Dyn2 in regulating podosome maturation and turnover. Despite numerous studies having been performed on podosome formation (Cervero, Wiesner et al., 2018, Labernadie, Bouissou et al., 2014, van den Dries, Bolomini-Vittori et al., 2014, van den Dries, Meddens et al., 2013), it largely remained unclear how the actin cores of podosomes retained their cylindrical shape under mechanical stress during protrusion. Based on our data, we hypothesize that Dyn2 binds and assembles around the actin cores to stiffen them, facilitates their assembly into the columnar architecture and, finally, triggers podosome turnover via GTP hydrolysis-induced disassembly (Fig. 8f) (Chuang et al., 2019). Therefore, we speculate that Dyn2 functions as a molecular girdle to maintain the structure of actin cores when podosomes encounter physical stress.

In contrast to our understanding of podosome formation, relatively little is known about how podosome turnover is regulated. To date, only myosin II, Supervillin and fascin have been reported to regulate podosome turnover (Bhuwania, Cornfine et al., 2012, Van Audenhove, Debeuf et al., 2015, van den Dries et al., 2013). Myosin II and Supervillin enable podosome turnover by increasing actomyosin contractility, whereas fascin facilitates podosome disassembly by inhibiting Arp2/3-mediated actin branching. Here, we have shown that GTP hydrolysis-induced Dyn2 disassembly is also involved in podosome turnover. These observations demonstrate that both formation and turnover of podosomes are tightly regulated in cells and that Dyn2 plays roles in both processes. Further studies are needed to better understand the multiple regulatory mechanisms controlling podosome turnover.

Although the function of Dyn2 at the presynaptic membrane has been well studied (Chung, Barylko et al., 2010, Hayashi, Raimondi et al., 2008, Newton, Kirchhausen et al., 2006), the role of Dyn2 at the postsynaptic membrane is largely unknown. Consistent with the function of synaptic podosomes in NMJ maturation (Proszynski & Sanes, 2013), we have shown here that postsynaptic expression of Dyn2 mutants with a defect in podosome turnover results in abnormal AChR cluster morphology in cultured myotubes and a disorganized pattern of spectrin distribution in *Drosophila* NMJs. Defects of postsynaptic spectrin and actin have been reported to disrupt SSR integrity, active zone spacing, glutamate receptor clustering, and electrophysiological activity (Blunk, Akbergenova et al., 2014, Pielage et al., 2006, Proszynski et al., 2009). Given that postsynaptic expression of Dyn2^A618T^ impaired the organization of postsynaptic spectrin, we reason that Dyn2^A618T^ expression causes a milder defect in *Drosophila* NMJs than spectrin knockdown.

CNM-associated Dyn2 mutations are hypermorphic alleles resulting from loss of autoinhibitory regulation, which promotes self-assembly (Faelber, Gao et al., 2013, Hohendahl, Roux et al., 2016). Hyper-assembled Dyn2 is hyperactive, resistant to GTP hydrolysis-induced disassembly, and presents enhanced membrane fission activity (Chin et al., 2015, Kenniston & Lemmon, 2010). Similar to observations of hyper-assembly of CNM-associated Dyn2 mutant protein on membrane, we show that Dyn2^A618T^ has stronger actin bundling ability, less sensitivity to GTP, and also extended podosome lifespan. Importantly, the phospho-mimetic Dyn2^Y597E^ mutant had a similar effect to Dyn2^A618T^, demonstrating that self-assembly is a critical switch of Dyn2 function in cells.

The structure of dynamin oligomers around membrane templates has been beautifully resolved; it binds to membrane via the PH domain and self-assembles into helixes through its stalk region (Antonny et al., 2016, Kong, Sochacki et al., 2018). Interestingly, our negative-stain TEM images revealed similar ring-like Dyn2 oligomers surrounding actin filaments. Akin to Dyn2 spirals on lipid templates, these Dyn2 oligomers on actin filaments are also responsive to GTP hydrolysis, which induces Dyn2 disassembly. Combined, these findings demonstrate that Dyn2 is a unique actin-binding protein that aligns and packs actin filaments together by forming ring-like oligomers around them. Recently, it was reported that the Dyn2 homolog in *Drosophila, shibire*, is also a multifilament actin-bundling protein, with *shibire* rings being located within the actin bundles (Zhang, Lee et al., 2020). Given that dynamin-actin bundling assays were conducted in non-physiological salt concentrations of 50 mM and 75 mM KCl in their and our experiments, respectively, we are conscious of the artificial conditions we both used. Therefore, parallel *in vivo* experiments, together with careful interpretation of *in vitro* reconstitution results, are critical to study the interplay between dynamin and cytoskeleton proteins (Shpetner & Vallee, 1989). Since our results demonstrated recruitment of Dyn2 to synaptic podosomes after formation of actin cores, we speculate that Dyn2 binds to actin filaments of podosome cores and assembles around them.

In summary, Dyn2 is an evolutionarily conserved actin bundler that regulates synaptic podosome maturation and turnover, where it participates in the development of NMJs. Many questions remain unanswered regarding exactly how Dyn2 binds and assembles on actin filaments, whether Dyn2 assembly at podosomes is regulated by other interacting protein, and if Dyn2 contributes to postsynaptic maturation in synapses other than NMJs. Our study highlights a distinct regulatory molecule involved in podosome turnover and the necessity for further studies on podosome kinetics and mechanical properties.

## Materials and Methods

### Cell culture

Mouse-derived C2C12 myoblasts (ATTC, CRL-1772) were cultured in growth medium containing DMEM supplemented with 2 mM L-glutamine, 1 mM sodium pyruvate, antibiotics, and 10% FBS (Gibco). To induce cell differentiation, growth medium was replaced with differentiation medium containing DMEM supplemented with 2 mM L-glutamine, 1 mM sodium pyruvate, antibiotics, and 2% horse serum (Gibco) at 90% confluency. This time-point was considered as day 0 of differentiation. c-SrcY527F transformed NIH3T3 cells were cultured in DMEM, 10% FBS and 200 μg/ml Hygromycin B as previously described (Pan, Chen et al., 2011).

For immunostaining and time-lapse imaging, coverslips, Permanox slides (#160005, Thermo Fisher Scientific), or glass-bottom dishes (#P35G-1.5-14-C, MatTek) were coated with 10 μg/ml of laminin (#23017-015, Invitrogen) and incubated overnight at 37 °C prior to seeding. The laminin solution was aspirated completely before plating cells. Cells were seeded at 80% confluency one day before differentiation. After differentiation, cells cultured on laminin-coated coverslips or Permanox slides were ready for immunostaining, and cells cultured on laminin-coated glass-bottom dishes were ready for time-lapse imaging.

### Transfection, lentiviral and adenoviral infection

For transfection, cells at 70% confluency were transfected with target DNA using Lipofectamine 2000 (#11668-027, Invitrogen), as recommended by the manufacturer. For Dyn2 knockdown experiments, lentiviruses with targeting shRNA sequences [5’-GCCCGCATCAATCGTATCTTT-3’ (#1) and 5’-GAGCTCCTTTGGCCATATTAA-3’ (#2)] were prepared and used. C2C12 myoblasts at 50% confluency were infected with viruses and selected with 2 μg/ml puromycin for three days. Cells surviving after selection were pooled together to reach 90% confluency for differentiation. For adenoviral infection, day 3-differentiated myotubes were infected with viruses overnight in the presence of differentiation medium containing 10 ng/ml tetracyclin. Constructs used in this study are listed in Supplementary Table 2.

### Cell and mouse TVA muscle staining

C2C12-differentiated myotubes were fixed with 4% paraformaldehyde at 37 °C for 2 min and continued fixation at room temperature for 5 min. After washing with PBS, cells were permeabilized in PBS plus 0.1% saponin and treated with PBS plus 2% BSA and 5% normal donkey serum for blocking. Cells were then stained with the indicated primary and secondary antibodies. Antibodies used in this study are listed in Supplementary Table 3.

The *transversus abdominis* (TVA) muscles were dissected from 4-week-old wild-type mice and fixed in 4% paraformaldehyde, as previously described (Au - Murray, Au - Gillingwater et al., 2014). To visualize NMJs in mouse TVA muscle, after being permeabilized in 2% Triton X-100 and blocked in 4% BSA plus 1% Triton in PBS, the TVA muscles were washed in PBS and stained using the indicated antibodies. After washing in PBS, all samples were mounted in Fluoromount-G (#0100-01, SouthernBiotech).

### Fluorescence microscopy

For fixed samples, images were collected using an LSM700 confocal microscope with a 63×, 1.35-NA oil-immersion objective (Carl Zeiss), an LSM780 confocal microscope (Carl Zeiss), or a TSC SP8 X STED 3X (Leica) system with a 100× oil objective 1.4 NA (STED microscopy), acquired with the excitation laser at 488 or 594 nm and the depletion laser at 592 or 660 nm using Hybrid Detector (Leica HyD).

For time-lapse microscopy, cells were seeded on laminin-coated glass-bottom dishes and transfected with target DNA. After 5 days of differentiation, the differentiation medium was replaced with imaging medium (phenol-red free DM with 20 mM HEPES, pH 7.4, 50 μg/ml ascorbic acid, and 10% FBS). Cells were imaged with a Zeiss inverted microscope Axio Observer Z1 at 37 °C.

To image reconstituted actin bundles, 1 μM F-actin solution (20% rhodamine-labeled) was applied to chambers loaded with acid-washed coverslips. Actin polymerization buffer containing 0.2 μM Dyn2 and 0.2 mM GTP was added afterward and imaged under a LSM780 confocal microscope (Carl Zeiss) at room temperature.

### Matrix degradation assay

To make FITC-gelatin-coated coverslips, acid-washed coverslips were first coated with 0.01% poly-D-lysine (#P7280, Sigma-Aldrich) for 1 h at room temperature and washed three times with PBS. We added 0.5% glutaraldehyde (#G5882, Sigma-Aldrich) to coverslips on ice for 15 min and washed with cold PBS. Pre-warmed coating solution composed of 0.1 mg/ml FITC-gelatin (#G-13187, Invitrogen) and 10 μg/ml laminin (#23017-015, Invitrogen) were added to the coverslips, which were then placed in the dark for 10 min at room temperature. After washing with PBS, 5 mg/ml NaBH4 (#452882, Sigma-Aldrich) was added for 15 min to inactivate residual glutaraldehyde. The FITC-gelatin-coated coverslips were washed with PBS and stored in 70% ethanol.

Day 3-differentiated C2C12 myotubes were plated on FITC-gelatin-coated coverslips with or without adenovirus induction. After 16 h, the cells were fixed, stained for F-actin and HA-Dyn2, and imaged by confocal microscopy. The thresholding feature in Metamorph software (Molecular Devices) was used to analyze matrix degradation areas and cell-containing areas.

### Fly stocks and genetics

Fly stocks and *GAL4* lines were obtained from the Bloomington *Drosophila* Stock Center and maintained on normal food medium. The parental strain ZH-51D was used to generate transgenic flies by injecting the *pUAST-based* constructs into *Drosophila* embryos, thereby integrating them into the attP (second chromosome) landing site.

### Dissecting and staining of larval body wall muscle

Third-instar larvae were dissected in Ca^2+^-free buffer (128 mM NaCl, 2 mM KCl, 4 mM MgCl, 5 mM HEPES, 35.5 mM sucrose and 5 mM EGTA) and fixed in either 4% formaldehyde for 20 min or Bouin’s fixative (Sigma-Aldrich) for 2 min, and then rinsed in 0.1 M phosphate buffer (pH 7.2) containing 0.2% Triton X-100. After blocking, samples were incubated overnight with indicated primary antibody at 4 °C. After further staining with secondary antibodies, samples were mounted in Fluoromount-G (#0100-01, SouthernBiotech).

### TEM of *Drosophila* NMJs

Larval fillets were dissected in calcium-free HL-3 medium (70 mM NaCl, 5 mM KCl, 10 mM MgCl_2_, 10 mM NaHCO_3_, 5 mM HEPES, 115 mM sucrose, 5 mM Trehalose, pH 7.2) at room temperature and were fixed for 12 h in 4% paraformaldehyde/1% glutaraldehyde/0.1 M cacodylic acid (pH 7.2) solution and then rinsed with 0.1 M cacodylic acid (pH 7.2) solution. They were subsequently fixed in 1% OsO_4_/0.1 M cacodylic acid solution at room temperature for 3 h. The samples were subjected to a series of dehydration steps, i.e., from 30% to 100% ethanol. After the 100% ethanol dehydration step, the samples were incubated in propylene, a mixture of propylene and resin, and then in pure resin. Lastly, they were embedded in 100% resin. The images of type Ib boutons were captured using a Tecnai G2 Spirit TWIN system (FEI Company) and a Gatan CCD Camera (794.10.BP2 MultiScan) at ≥4,400× magnification. We identified type Ib boutons by the multiple layers of subsynaptic reticulum, and the size and layers of SSR of type Ib boutons were measured using Image J (NIH) accordingly to previous reports (Budnik, Koh et al., 1996, Lee & Schwarz, 2016). In brief, SSR thickness was measured in ImageJ as follows: (1) the center of mass was determined by drawing a region of interest around the periphery of a bouton; (2) four lines were drawn 90 degrees apart from one another emanating from the center of mass; and (3) SSR thickness was then determined based on the average length between SSR edge and bouton edge along each line. SSR layers were determined by the plot profile feature in ImageJ across the segment of SSR on each line.

### Electrophysiology

Evoked excitatory junctional potential (EJP) was recorded as previously described (Peng, Lin et al., 2019). Briefly, third instar larvae were dissected in calcium-free HL3 buffer at room temperature, followed by incubation in 0.5 mM Ca^2+^ HL3 solution for 5–10 min prior to recording. The mean resistance value for the recording electrode was ~40 MΩ when 3 M KCl solution was used as the electrode solution. All records were obtained from muscle 6 in the A3 hemisegment. Resting membrane potentials of muscles were held at less than −60 mV. EJPs were amplified using an Axoclamp 900A amplifier (Axon Instruments) under bridge mode and filtered at 10 kHz. EJPs were analyzed using pClamp 10.6 software (Axon Instruments). Averaged EJP amplitude was calculated from the amplitudes of 80 EJPs in one consecutive recording. Miniature EJP recordings were performed in HL3 solution containing 0.5 mM Ca^2+^ and 5 μM tetradotoxin (TTX) and also analyzed using pClamp 10.6 software.

### Image analysis

Immunostained images were analyzed in Metamorph (Molecular Devices) and ZEN (Carl Zeiss) software. Cell area and *Drosophila* NMJ area were selected manually and the area and signal intensity were measured in Metamorph. Matrix degradation area as well as GluRIIA and AChR cluster area were selected using threshold features in Metamorph and quantified automatically using the same software. Podosome diameter and height were analyzed manually in xz orthogonal view in ZEN.

### F-actin bundle sedimentation assay

Dynamin proteins were expressed in Sf9 cells transiently transfected with various constructs and purified as previously described, then snap-frozen in buffer containing 20 mM HEPES, 150 mM KCl, 1 mM EGTA, 1 mM DTT and 10% glycerol (Liu, Neumann et al., 2011). The actin bundling assay was performed as described previously (Chuang et al., 2019, Lin, Chuang et al., 2019). Briefly, 10 μM purified rabbit skeletal muscle actin (#AKL99-C, Cytoskeleton) was diluted in general actin buffer (5 mM Tris pH 7.4, 0.2 mM ATP, 0.2 mM CaCl2, and 0.5 mM DTT) and incubated at 4 °C for 1 h and then centrifuged at 20,000 *g* for 15 min to remove aggregated proteins. The G-actin was polymerized in actin polymerization buffer comprising 2.5 mM Tris pH 7.4, 0.1 mM CaCl2, 0.25 mM DTT, 50 mM KCl, 2 mM MgCl2, and 1 mM ATP for 1 h at room temperature to generate actin filaments. To reconstitute branched actin, 160 nM WASP VCA domain protein (#VCG03, Cytoskeleton) and 60 nM Arp2/3 protein complex (#RP01P, Cytoskeleton) were also added to the actin polymerization buffer. To generate Dyn2-actin bundles, 5 μM polymerized F-actin was incubated with the indicated concentrations of Dyn2, resulting in a final KCl concentration of 75 mM. After 30-min incubation at room temperature, the mixture was centrifuged at 14,000 *g* for 20 min at room temperature. Protein in supernatants and pellets was solubilized in SDS sample buffer and subjected to SDS-PAGE. Proteins were visualized by Coomassie blue staining, and band intensities were quantified using ImageJ.

### Transmission EM

To visualize actin bundles, 5 μM filamentous actin was incubated with or without 1 μM Dyn2 at room temperature for 30 min. The mixture was then diluted 2-fold and adsorbed onto carbon-coated grids and stained with 2% uranyl acetate. Images were collected using a Hitachi H-7650 electron microscope at 75 kV and a nominal magnification of 120,000. To image Dyn2-actin bundles upon addition of GTP, the actin mixture was placed on parafilm before GTP addition. GTP was added to the actin mixture with a final concentration of 1 mM. The carbon-coated grids were placed on top of the mixture one min before being subjected to 2 % uranyl acetate staining. Negative-stained samples of actin bundles in the presence of other nuceotides were prepared and captured by TEM as described above.

### Statistical analysis

Quantitative data are expressed as mean ± SD of at least three independent experiments. All data were analyzed with one-way ANOVA or Student’s *t* test. Statistical significance was defined using GraphPad Prism 8.0. *P* < 0.05 was considered statistically significant, indicated as *, *P* < 0.05; **, *P* < 0.01; or ***, *P* < 0.001.

## Acknowledgments

We thank the staff of the imaging core at First Core Labs, National Taiwan University (NTU) as well as the EM core in NTU, the EM core in the Institute of Molecular Biology at Academia Sinica, and the Taiwan Protein Project of Academia Sinica for their technical support. We are grateful to Dr. Tomasz Proszynski (Nenki Institute of Experimental Biology) for sharing his protocol for visualizing AChR clusters in cultured myotubes. We thank Dr. Chun-Fang Huang (National Laboratory Animal Center) for helping with mouse NMJ isolation and staining. We also thank Prof. Allen Liu (University of Michigan) and Chun-Liang Pan (NTU) for helpful comments and critical reading of this paper. This work was supported by Ministry of Science and Technology grants MOST 107-2628-B-002-008 and MOST 108-3017-F-002-004, National Taiwan University grant NTU-109L7808 to Y.W. Liu. Part of this work was supported by Taiwan Protein Project grant AS-KPQ-105-TPP.

## Author contributions

All authors participated in experimental design. SSL, TLH, GGL, TNL, HCL, CWC, HYW, CKY and YWL performed experiments. SSL and TLH analyzed data, and SSL, CKY, and YWL wrote the manuscript. CKY and YWL supervised the project.

